# Excess deaths associated with the chikungunya epidemic of 2014 in Jamaica were higher among children under 5 and over 40 years of age, an analysis based on official data

**DOI:** 10.1101/227579

**Authors:** André Ricardo Ribas Freitas, Maria Rita Donalisio

## Abstract

We assessed the excess of all causes of mortality by age groups during the chikungunya epidemics in Jamaica, 2014. Excess mortality was estimated by subtracting deaths observed in 2014 from that expected based on the average mortality rate of 2012-2013, with confidence interval of 99%.

Overall mortality 91.9 / 100,000 population, 2,499 additional deaths than expected coincided with the peak of the epidemic, there was a strong correlation between the monthly incidence and the excess of deaths (Spearman Rho = 0.939; p <0.005). No other significant epidemiological phenomenon occurred on that island that could explain this increase in mortality. Thus, we suggest that mortality associated with chikungunya is underestimated in Jamaica, as in other countries.

The excess of deaths could be a strategic tool for the epidemiological surveillance of chikungunya as it has already been used in influenza and respiratory syncytial.

## INTRODUCTION

Chikungunya is an emerging arbovirus first identified in the Americas in late 2013. On December 9th, 2013, the Pan American Health Organization[1] issued an alert on the transmission of chikungunya in the Caribbean and on the risk of geographic expansion. By the end of 2014, the transmission of chikungunya was already confirmed in most of the countries bathed by the Caribbean Sea, totaling 964,341 cases and 194 deaths[2].

In Jamaica the first cases of chikungunya were reported in May 2014, and confirmation of autochthonous transmission occurred on August 5th, 2014[3]. In October of the same year, the epidemiological situation worsened, prompting the Jamaican government to declare emergency [4], although no cases of death by chikungunya were reported by Jamaica to PAHO[1].

Since the re-emergence of the chikungunya in the first decade of the 21st century many studies have been published on severe cases and deaths caused by chikungunya[5–9]. However, studies in some Indian Ocean islands and continental India suggest that in some situations, the passive surveillance system may not recognize many of the deaths associated with chikungunya, but an excess of deaths during the epidemic period can be identified[10–12]. The objective of this study was to evaluate the mortality associated with the chikungunya epidemic of 2014 in Jamaica.

## METHODOLOGY

Jamaica is a tropical island (Aw, Köppen climate classification) with 2,720,554 inhabitants in the Caribbean Sea whose capital, Kingston, is located at latitude 17° 59′ N and longitude 76° 48′ W.

We obtained chikungunya incidence data from the Jamaica Open Site Portal [3], population data on the website [13] and mortality data were obtained from the site[14].

Monthly mortality rates were calculated for the years 2012 to 2014. The number of deaths expected for each month of 2014 was estimated using the monthly average observed in the years 2012 and 2013, projecting for the expected population of Jamaica in 2014. These years (2012 and 2013) were chosen to avoid changes in the population profile altering the results. The maximum threshold of deaths expected in 2014 was the upper limit of the 99% confidence interval. The mortality rate by age group between the years of 2012 and 2014 was also calculated, excluding deaths from external causes. The number of deaths expected by age group for the year 2014 was calculated using the average mortality by age group for the years 2012 and 2013 and the population estimated by age group. We define excess deaths as the difference between the observed and the expected.

## RESULTS

There was a significant increase in overall mortality during the year 2014, this increase was most significant during the chikungunya epidemic period, being above the upper limit of the 99% confidence interval (Figure 1) coinciding with the epidemic peak.

**Figure 1.**
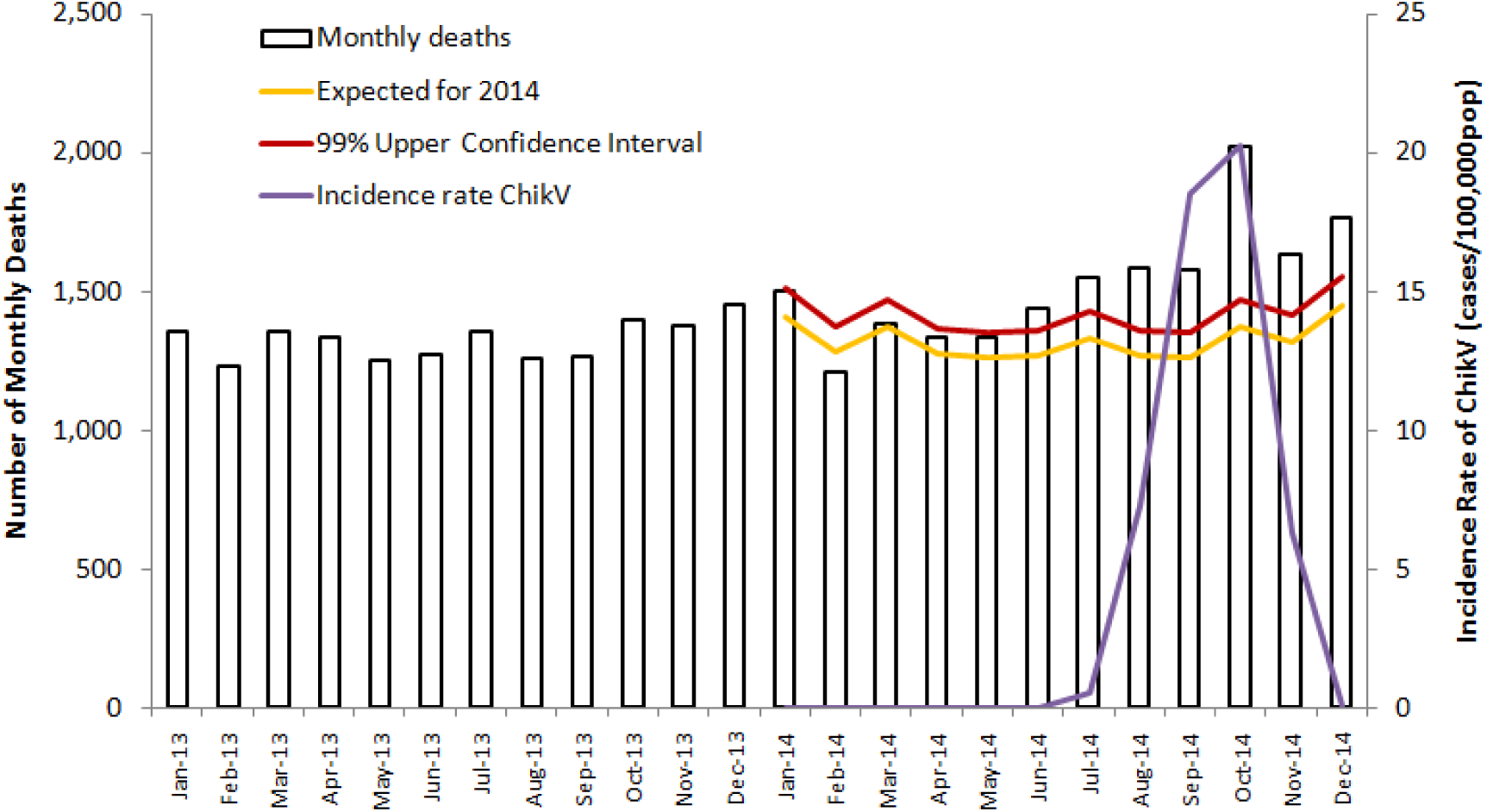
All causes monthly mortality (Jamaica, 2013-2014), expected deaths and upper 99% confidence interval (2014) and chikungunya incidence rate (cases/100,000 population).

There was a strong correlation between the monthly incidence coefficient of chikungunya and the excess of deaths during the epidemic period (Spearman Rho = 0.939; p <0.005). Overall mortality increased by 91.9 per 100,000 population, 2,499 more deaths than expected for the year 2014, above the average of the two years prior to the epidemic. The excess mortality was greater among the elderly, but there was also an increase of deaths above the 99% confidence interval in the age groups: up to 4 years, 20 to 29 years and in all age groups over 40 years (Table 1). The excess mortality rate in the 50-59 age group was 104.6 / 100,000 population (1 in 1,000) and among those over 70 years, 998.6 / 100,000population, close to one death for every 100 individual of this age group.

**Table 1.** Population by age group, expected, observed, excess deaths and excess mortality rate (deaths/100,000 population, Jamaica, 2014)

|  | Population | Expected Deaths for 2014 | UPPER 99% Confidence Interval | Observed Deaths 2014 | Excess Deaths | Excess Mortality Rate (deaths/100,000) |
| --- | --- | --- | --- | --- | --- | --- |
| 0 to 4 years | 199,170 | 694 | 742 | **<u>758</u>** | **<u>64</u>** | **<u>32.3</u>** |
| 5 to 9 years | 217,065 | 42 | 53 | 33 | -9 | -4.0 |
| 10 to 19 years | 523,010 | 117 | 136 | 132 | 15 | 2.9 |
| 20 to 29 years | 483,103 | 220 | 247 | **<u>274</u>** | **<u>54</u>** | **<u>11.1</u>** |
| 30 to 39 years | 376,788 | 491 | 531 | 531 | 40 | 10.7 |
| 40 to 49 years | 343,682 | 939 | 994 | **<u>1,101</u>** | **<u>162</u>** | **<u>47.3</u>** |
| 50 to 59 years | 257,205 | 1,500 | 1,570 | **<u>1,769</u>** | **<u>269</u>** | **<u>104.6</u>** |
| 60 to 69 years | 161,514 | 2,229 | 2,315 | **<u>2,544</u>** | **<u>315</u>** | **<u>194.9</u>** |
| 70 and more years | 159,017 | 8,916 | 9,087 | **<u>10,504</u>** | **<u>1,588</u>** | **<u>998.6</u>** |
| Total | 2,720,554 | 15,147 | 15,676 | **<u>17,646</u>** | **<u>2,499</u>** | **<u>91.9</u>** |

**Observed mortality rate above the 99% confidence interval was bold and underlined.**

## DISCUSSION

Several studies conducted over the past decade have shown an increase in overall mortality during chikungunya epidemics in the Indian Ocean[10–12, 15]. In the Reunion Islands most of the deaths were laboratory confirmed[6, 7, 9], and in some cases deaths by chikungunya were attested as a basic cause or as contributing cause some of them without mention of laboratory confirmation [5]. In the Réunion Islands (2006) epidemic, overseas French territory, there was a robust diagnostic support, resulting perhaps in the similarity in the number of deaths by chikungunya found in the death certificates and the estimates by excess of deaths. In contrast, in Mauritius and Nicobar, the increase in mortality was not accompanied by reports of chikungunya deaths, although there was a significant increase in overall mortality during the epidemic[10, 12, 16]. In Ahmedabad (India, 2006), there was a significant increase in overall mortality equivalent to 2,944 deaths during the epidemic. Despite scientific publications with laboratory confirmations of dozens of severe cases and deaths by chikungunya[8], incredibly there was no report of deaths by Indian public health authorities[11].

Since chikungunya began to be transmitted in the Americas, several authors have demonstrated the occurrence of laboratory confirmed cases of death[17–21] (Evans-Gilbert, 2017, Hotz, 2015, Sá, 2017, Rollé, 2016, Mendoza, 2015;, 2016). These deaths include patients at various ages, newborns, children, young and old adults, previously healthy people and patients with comorbidities. The main complications leading to death were myocarditis, septic shock, encephalitis, and respiratory complications [17–23]

The present study suggested a strong temporal correlation between general mortality and the chikungunya epidemic occurring in Jamaica. The age groups with the greatest increase in mortality during the epidemic period were the same identified in Reunion Island as chikungnuya deaths[5] and in Brazil through excess mortality[24].

There was no extreme weather phenomenon in Jamaica in 2014[25], neither a significant circulation of dengue virus or other arboviruses on that island in 2014[26], there was also no intense circulation of influenza virus[27], which is known to lead to an increase in overall mortality even in tropical countries[28]. Studies on excess of mortality in Brazil have found a number of deaths above that identified by passive, conventional epidemiological surveillance[24, 29].

The excess mortality found in Jamaica was 2499 deaths, more than 10 times the total deaths reported by all the countries of the Americas in the year 2014 for PAHO. It corresponds to a mortality rate of 91.9 / 100,000 inhabitants, almost twice the rate found in Pernambuco (47.9 / 100,000 inhabitants), the most affected state in Brazil by the chikungunya in 2016[24], and almost three times that found in the Reunion Islands in 2006 (33.8 / 100,000 inhabitants)[15].

This is an ecological time series study with limitations, however, we believe that our results corroborates the hypothesis that the chikungunya virus is contributing significantly to the increase in the number of deaths in Jamaica, and maybe in other countries in America in 2014, as well as the observed in the Indian Ocean and in the Americas since 2014.

We believe that mortality associated with chikungunya has been underestimated, this may be due to the poor access to diagnostic methods, but also may be associated with the difficulty of recognizing the severe forms of the disease [30]. The fact that most of the deaths occur in the elderly[5], associated to the fact that the clinical picture in this age group seems to be different from the one presented by younger patients[31], may lead the attending physician to have difficulty in establishing the relationship between the deaths and infection by chikungunya. In fact, a recently published study found that 42.7% of the elderly attended in a hospital in Martinique with laboratory-confirmed chikungunya did not present a typical clinical picture due to absence of fever, absence of joint pain or both[31]. More studies on clinical outcomes and prognostic of chikungunya infection should be developed to better understand the causes of chikungunya-related deaths. These studies may lead to changes in clinical management strategies in situations of epidemics as well as simultaneous circulation of more than one arbovirus[32].

Correct assessment of the impact of chikungunya as the cause of death is an important step in scaling up the burden of the disease. We propose that the evaluation of excess of deaths could be a strategic tool for epidemiological surveillance to monitor the mortality associated with chikungunya as it is already used for surveillance influenza, respiratory syncytial virus and extreme climatic phenomena [33–35]

